# Dietary zinc restriction induces irreversible allodynia in adult mice

**DOI:** 10.1101/2025.06.02.657430

**Authors:** Daiane Oliveira Matias, Bruna Lima Roedel dos Santos, Aline França Martins, Larissa Poblete de Souza, Leandro Miranda-Alves, Luiz Eduardo de Macedo Cardoso, Ana Luísa Palhares de Miranda, Luís Maurício T. R. Lima

## Abstract

**Introduction:** Neuropathic pain is a debilitating condition highly prevalent worldwide. One of the main factors related to its development is poor nutrition and micronutrient deficiency in the diet. Essential metals, especially divalent ions as zinc, play crucial roles in the neurobiology of the nervous system, including pain signaling and transmission mechanisms, and metabolism control. The present study aimed to understand the impact of zinc in neuropathic pain, evaluating whether restriction and direct repletion of a dietary zinc could change nociceptive behavior and metabolic parameters in adult mice.

**Methods:** Adult male *Swiss* mice received a zinc-restricted diet for eight weeks. The repletion group received the restricted diet for four weeks followed by normal zinc diet for another four weeks. Mechanical and heat thermal pain sensitivity were assessed using the von Frey filaments and Hargreaves tests, respectively.

**Results:** Zinc restriction resulted in decreased body weight gain and led to an increased mechanical and thermal sensitivity to heat. Dietary zinc repletion reversed the thermal allodynia and increased weight gain, abdominal adipose tissue, and liver weight compared to a normal zinc diet. No changes were observed concerning food and water intake, glycemic profile, pancreatic morphology and plasma amylin,

**Conclusions:** The reduction in the bioavailability of dietary zinc promotes metabolic and nociceptive changes in adult mice, inducing allodynia characteristic of neuropathic pain.

## 1. Introduction

The new definition and classification of chronic pain proposed by IASP was included for the first time in the International Classification of Diseases (ICD-11) of the World Health Organization. Thus, chronic pain is defined as “pain that persists or recurs for more than three months”, categorized into primary and secondary chronic pain. Primary chronic pain is multifactorial: biological, psychological and social, and secondary chronic pain is related to various health conditions that have pain as part of their outcome, such as painful neuropathies (Treede et al. 2019; Raja et al. 2020). Metabolic diseases, such as diabetes, nutritional conditions, vasculitis, infections and genetic factors can trigger neuropathic pain (Mills et al. 2019; Cole and Florez 2020).

Diabetes mellitus defines a group of disorders of insulin and amylin physiology (American Diabetes Association 2022). Diabetes is increasing worldwide for both type 1 (T1DM) and type 2 (T2DM) (Ong et al. 2023), indicating prevalence of environmental components in the disease process, which may include pollutants, synthetic chemical endocrine disruptors and nutrition. T1DM and T2DM degeneration of the β-cell health occurs by multiple mechanisms and in different time scales, although residual insulin production is found in T1DM adults (Harsunen et al. 2023).

Diabetes has multiple extra-pancreatic manifestations, occurring systemically, both as cause and as the result of the disease process, and multiple comorbidities (including metabolic syndrome components) and complications (including neuropathy, nephropathy, retinopathy, microvascular) (Lima 2017). Diabetic neuropathy (DN) is one of the most prevalent and debilitating, which, according to the Diabetes Control and Complications Trial (DCC), affects approximately 50% of patients over time. DN is characterized by the progressive loss of sensory function, resulting in increased mechanical, thermal sensitivity and symptoms related to chronic pain, that severely impact quality of life (Cole and Florez 2020).

Epidemiological evidence has suggested that micronutrient deficiency may participate on the onset of diabetes (Kaur and Henry 2014; Mente et al. 2017), including zinc (Wijesekara et al. 2009; Miao et al. 2013; Vashum et al. 2013; Carvalho et al. 2017). Mice lacking the zinc transporter (ZnT8 -/-) behaved most likely normal mice in physiology compared to the background strain, although lacking non-crystalline insulin in secretory granule and becoming glucose intolerant upon high-fat diet feeding (Lemaire et al. 2009). While a complete impairment of zinc transport to secretory granule disrupts intra-organelle interactions, dietary zinc restriction in normal weaning mice did not alter insulin crystallinity in secretory granules, although resulting in apoptosis and degeneration of the endocrine pancreas (Sisnande et al. 2020) and increased allodynia (Lima et al. 2023). The degenerative manifestations found in pancreatic islets of zinc-restricted animals are compatible to non-autoimmune T1DM (T1b) due to micronutrient imbalance, motivating further investigation of the consequences of essential micronutrient deficiency on the onset of diabetes.

The role of nutrition in the development and prevention of chronic pain is not yet fully understood. However, proper nutritional management is crucial for overall human health and can be beneficial for pain management. This is particularly important because good nutrition is associated with improvements in factors related to diseases like diabetes, which often involve chronic pain as a symptom.

Elma and collaborators conducted a systematic review of 24 pre-clinical trials on dietary modulation. Their findings revealed that diet composition can influence pain perception, having either analgesic or pro-nociceptive effects. Dietary patterns high in saturated fat, monounsaturated fat, and refined carbohydrates were found to lower the sensitivity threshold to nociceptive pain, leading to mechanical allodynia and heat hyperalgesia in inflammatory pain. Conversely, a diet rich in anti-inflammatory ingredients, such as omega-3 fatty acids, resveratrol, curcumin, sulforaphane, and ginseng, as well as a calorie-restricted diet, promoted recovery from both mechanical allodynia and heat hyperalgesia in chronic inflammatory pain (Elma et al. 2022). The low supply and dyshomeostasis of minerals, especially zinc, an essential divalent metal, may be directly related to the development and worsening of chronic pain (Philpot and Johnson 2019; Pagliai et al. 2020).

We have previously investigated the susceptibility of young mice to a zinc restriction since weaning and the pain response pattern. We demonstrated that a low zinc diet favors the development of mechanical allodynia and alters the expression of inflammatory markers such as TNF-α and the main enzyme of the antioxidant system that is structurally formed by copper and zinc, SOD-1, confirming the importance of these micronutrients for different types of pain (Lima et al. 2023). However, the consequences of zinc restriction on endocrine pancreas and nociceptive pain in adult animals, and whether zinc restriction complications are reversible, are unknown. In this study we report the investigation of the effect of zinc depletion and repletion in time-course of metabolism, insulin production and allodynia in adult mice.

## 2. Material and Methods

### Material

Purified rodent diets were formulated as described in details previously (Sisnande *et al*. 2020), based on the AIN93G (Reeves et al. 1993). The control diet was formulated with the AIN93G mineral mix (AIN93G-MX-control) and the zinc-deficient diet was formulated with a mineral mix without zinc carbonate (AIN93mG-MX-ZnDef). All reagents were of analytical grade and used as received.

### Animals

Swiss male mice were housed at vivarium at 22 ± 2°C, 60-80 % humidity, from the Faculty of Pharmacy (UFRJ), with 12-hour dark-light cycle. Animals had free access to water and food (Nuvilab CR1; Nuvital Nutrientes S/A, Quimtia Brasil; until beginning dietary intervention). The present study was conducted with approval by the Animal Care Committee of Federal University of Rio de Janeiro (CEUA-UFRJ) under protocol CCS-UFRJ-086-2021.

### Experimental Design

The experimental design is summarized in **Figure 1**.

**Figure 1.**
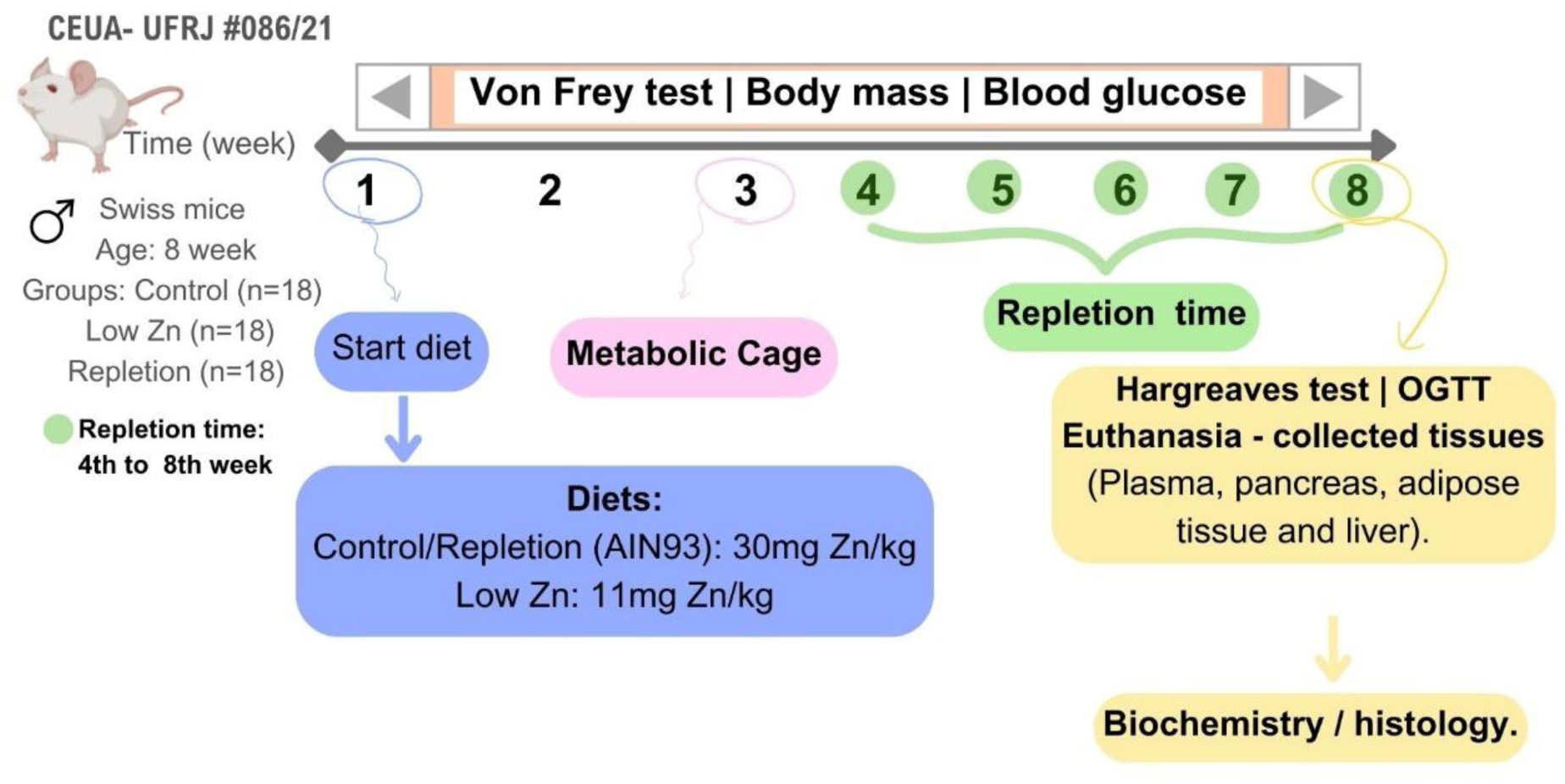
Study design. Swiss male mice were randomized into three experimental groups (Control, Low Zn, and Repletion; n=18 per group). Animals were tested weekly for body mass, capillary glycemia and mechanical allodynia. Metabolic cage was conducted in the 3^rd^ week of intervention, and in the 4^th^ week there was zinc replacement for the repletion group. In the 8^th^ week, the Hargreaves test was performed, followed by euthanasia for collection and weighing of organs and tissues.

Eight weeks old mice were randomized into the three arms of the study, receiving different purified rodent diets as described in details previously (Sisnande *et al*. 2020), named control (normal zinc content), restriction (low zinc) and repletion (normal zinc content after four weeks under zinc restriction) groups with 8 to 10 animals per group.

Animals were evaluated weekly for their body mass, capillary glycemia and mechanical (Von Frey test) allodynia. Hargreaves’s thermal sensibility test (heat allodynia) was performed at the eighth week of dietary.

Evaluation of the food and water consumption, feces and urine production were accessed by metabolic cage at third week of intervention, before the repletion.

The capillary glycemia was measured in the light cycle with a digital glucometer (AccuChek Active^®^, Roche^®^) from the tail tip after a small incision (about 1 mm) (Han et al. 2008).

At termination, biological specimens were collected, prepared for histological analysis and/or stored at – 80 °C for further analysis.

### Oral Glucose Tolerance Test (OGTT)

Oral glucose tolerance test was performed by standard protocol by an oral glucose load of 2 g/kg body weight (Pedro et al. 2020). The baseline glycemia was measured after 6h fasting, followed by gavage with a freshly prepared glucose solution (30 % w/v). Further measurements were performed at 15 min, 30 min, 60 min, 90 min and 120 min after oral glucose load.

### Metabolic Cage

Measurements of food and water consumption, feces and urine production, were carried out using a metabolic cage (Cat # 3600M021; Tecniplast, Italy) for 12 h during the dark cycle (22 ± 2°C; 60-80 % relative humidity). The animals were kept individually, and consumption was calculated as the difference between the initial available and the remaining quantities.

### Mechanical Allodynia - Von Frey Test

The evaluation of mechanical allodynia was conducted using the Von Frey test with the “up and down” method (Decosterd and Woolf 2000). The animals were individually placed in acrylic boxes (9 x 7 x 11 cm^3^) over a metallic surface with adequate mesh allowing access to animal’s paws using calibrated filaments (0.008 g to 2.0 g; Bioseb Lab Instr, Cat #BIO-VF-M) for mechanical stimuli. After adaptation to the box, the animal’s sensibility to varying strength filaments were measured, starting with 0.6 g for 5 times with 60 sec intervals between the stimuli, migrating to higher or lower strengths, according to the response until convergence of response (three repeated withdrawal values of five stimuli). The mechanical threshold was defined as the last strength filament that originated the highest number of paw withdrawal responses. Mechanical sensitivity was measured weekly.

### Heat Thermal Allodynia - Hargreaves test

The thermal sensitivity was evaluated using the Hargreaves test (Hargreaves et al. 1988). Animals were allocated in a glass surface for 30 min of acclimation. After that, a beam of radiant heat is applied to the center of the hind paw, through a mobile heating source (high intensity light) positioned under the glass floor. The heating stimulus was interrupted upon paw withdrawn indicating nociceptive response and the time between light turning on and off was considered as the latency time (in seconds). To avoid plantar tissue damage, the upper cut-off limit was 20 sec. The hind paw was evaluated three times, respecting an interval of five minutes, and the result expressed as the average of readings.

### Histological analysis

After euthanasia, the mouse pancreases were isolated, washed with PBS (137 mM NaCl, 2.7 mM KCl, 8 mM Na_2_HPO_4_ and 2 mM KH_2_PO_4_, pH 7.4) and fixed with buffered 4 % paraformaldehyde (pH 7.4). Three micron-thick slices from tissues embedded in paraffin were obtained for further staining and evaluation by optical microscopy (Olympus model BX53F, Olympus Optical, Co. Japan; digital camera DP- 72; Imaging Software CellSens).

### Histochemical Analysis

The area of the Langerhans islet was measured using Image J Fiji software (Schindelin *et al*. 2012) from the Hematoxylin-eosin (HE) staining.

The presence of amyloid deposits in the islets was investigated by staining the pancreatic tissue sections with Congo red (Merck, Cat #1.01340, Burlington, MA, USA) as described elsewhere (Westermark et al. 1999). Sections were then observed on a light microscope (Olympus, BX51, Tokyo, Japan) under bright field or polarized light, and images were acquired using a digital camera (Olympus, DP71, Tokyo, Japan). Positivity of the reaction for amyloid was checked against appropriate controls, which yielded a conspicuous red staining under bright field and a characteristic light-green birefringence under polarized light.

### Immunohistochemical Analysis

Pancreatic tissue was collected and fixed in 4% paraformaldehyde. Paraffin sections were dewaxed in xylol and hydrated in ethanol and distilled water. For antigen retrieval, the sections were incubated at 95°C for 35 min in 10 mM citrate buffer pH 6.0. Thereafter, the slices were rinsed in PBS (pH 7.4) and the endogenous peroxidase was blocked with 30% hydrogen peroxide in methanol for 20 min. Subsequently, blocking was performed with PBS 5% bovine serum albumin, 0.25% Triton X-100, 0.025% Tween-20 and 0.1% gelatin. The sections were incubated overnight in a humid chamber at 4°C with a primary anti-insulin antibody (insulin antibody, cat #4590, Cell Signaling) at a 1:100 dilution for the peroxidase assay. After washing with PBS pH 7.4 containing 0.25% Tween X-100, the tissues were incubated for 1 hour at room temperature with Histofine® Simple Stain MAX PO (cat #414341F, Nichirei Biosciences Inc. Tokyo. Japan) and counterstained with Harris’s hematoxylin.

### Statistical Analysis

All results are presented as mean ± standard error of the mean (SEM). Statistical analyses were performed using GraphPad Prism ver 8.2.0 (GraphPad Software). Two-way ANOVA, One-way ANOVA Tukey’s multiple comparisons test, or *t*-test were used depending on the experiment. Bonferroni and Sidak’s post-tests were used for multiple comparison analysis. A p-value of < 0.05 was considered significant. To indicate the magnitude, we used * p < 0.05, ** p < 0.01, *** p < 0.001. # Low Zn vs Repletion groups comparisons.

## 3. Results

### Metabolic impact of zinc restriction and repletion

Adult male mice divided into three groups were maintained on a normal zinc diet (*control group*) or low zinc diet (two groups: *restricted* and *repletion* group) for four weeks, after which the *repletion* group switched to the normal zinc diet, and all groups were maintained under experimentation for further four weeks.

The three groups gained body weight during the first four weeks (**Figure 2A**), although the two groups under zinc restriction showed a lower body weight gain rate. From the fifth to the eighth week of intervention, the *control* group reaches a plateau, while the *depletion* group showed a progressive decrease and the *repletion* group a significant gain of body weight (p = 0.0017), even higher than control group (**Figure 2B**). The group under zinc restriction showed a moderate loss in adipose tissue after eight weeks while the repletion group showed a significant increase in adipose tissue compared to the control (p = 0.0046) and Low Zn (p = 0.0002) (**Figure 2C**). Food (**Figure 3A**) and water (**Figure 3B**) intake, urine (**Figure 3C**) and feces (**Figure 3D**) production were not significantly affected by zinc restriction.

**Figure 2.**
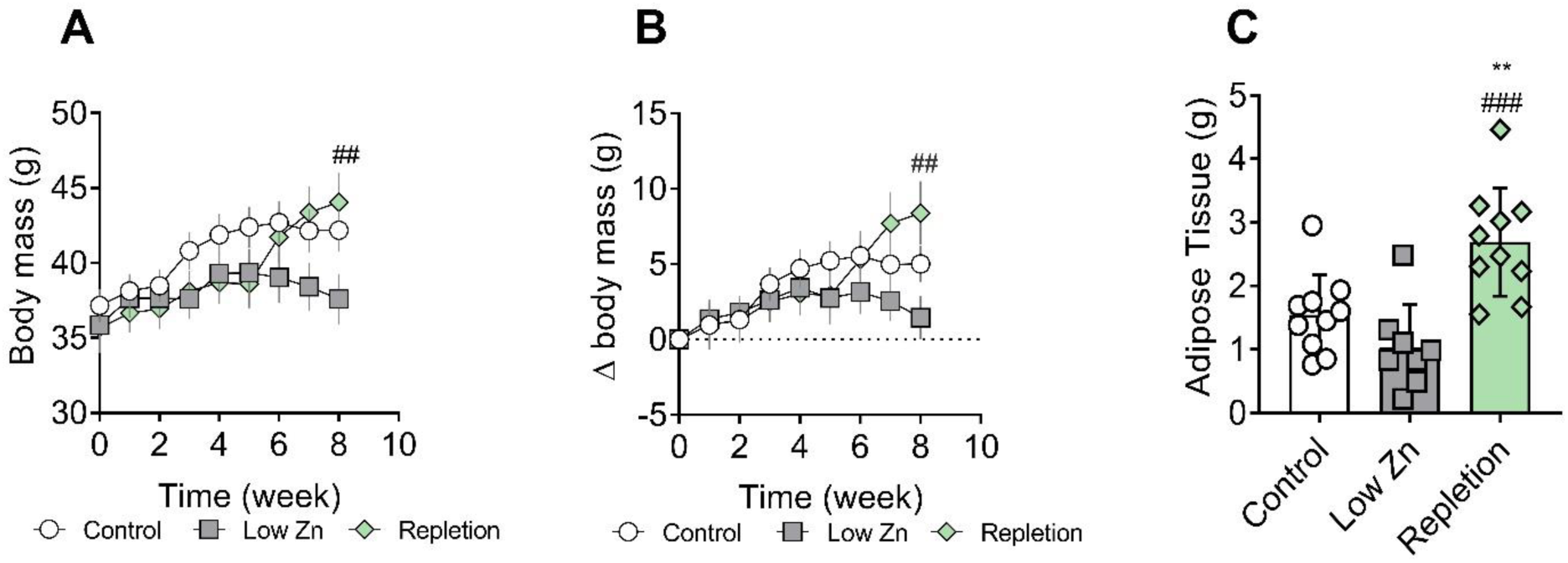
Effect of dietary zinc on body weight and abdominal adipose tissue in adult mice. **A)** body weight trajectory of control (**circles**), restriction (**squares**) and repletion (**green hexagon**) groups. **B)** Body weight gain curve. **C)** Abdominal adipose tissue. Repletion was initiated in the 4th week of intervention. Data are expressed as mean ± SEM (n=7-10). # p < 0.05 and ## p < 0.01 compared to Low Zn, Two-way ANOVA followed by Sidak’s multiple comparison post-test (**A** and **B**). ### p < 0.001 compared to Low Zn and ** p < 0.01 compared to control, One-way ANOVA test followed by Tukey’s post-test (**C**).

**Figure 3.**
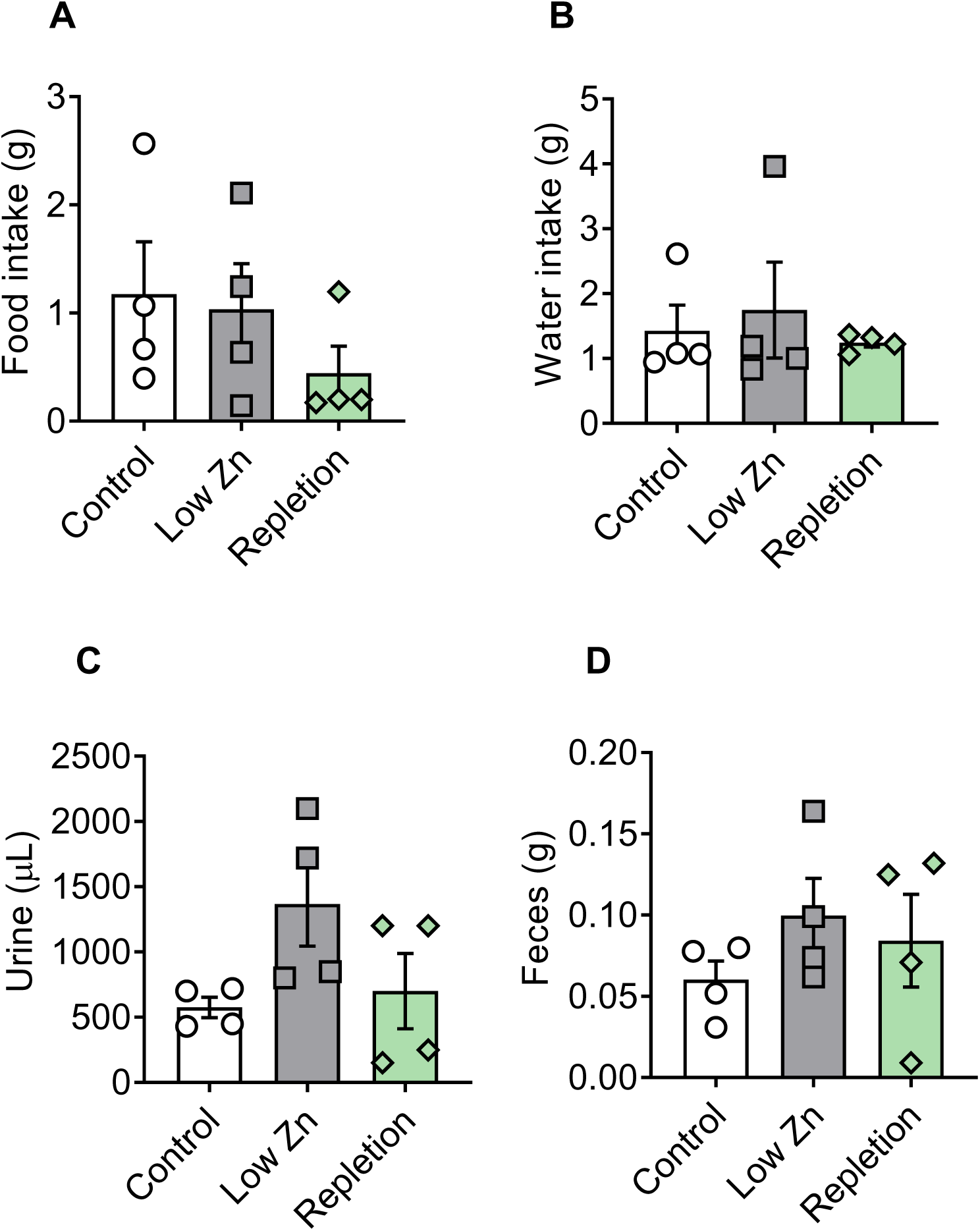
Effect of dietary zinc on nutrient metabolism. The animals that consumed a control, restricted or repletion zinc diet were evaluated regarding food consumption (**A**), water consumption (**B**), urine (**C**) and feces (**D**) production, at the 3rd week of intervention. Results are expressed as mean ± SEM (n=4). No significant differences were observed (p>0.05).

We did not observe significant differences in non-fasting glycemia (**Figure 4A**) and oral glucose tolerance (**Figure 4B**) throughout the intervention, while liver mass was significantly smaller in the restriction group (p = 0.0323) and higher in the repletion group compared to the control (p = 0.0223) and Low Zn (p = 0.0001) groups (**Figure 4C**). Circulating amylin, which is related to satiety and gastric emptying, it was not different between the groups (**Figure 4D**).

**Figure 4.**
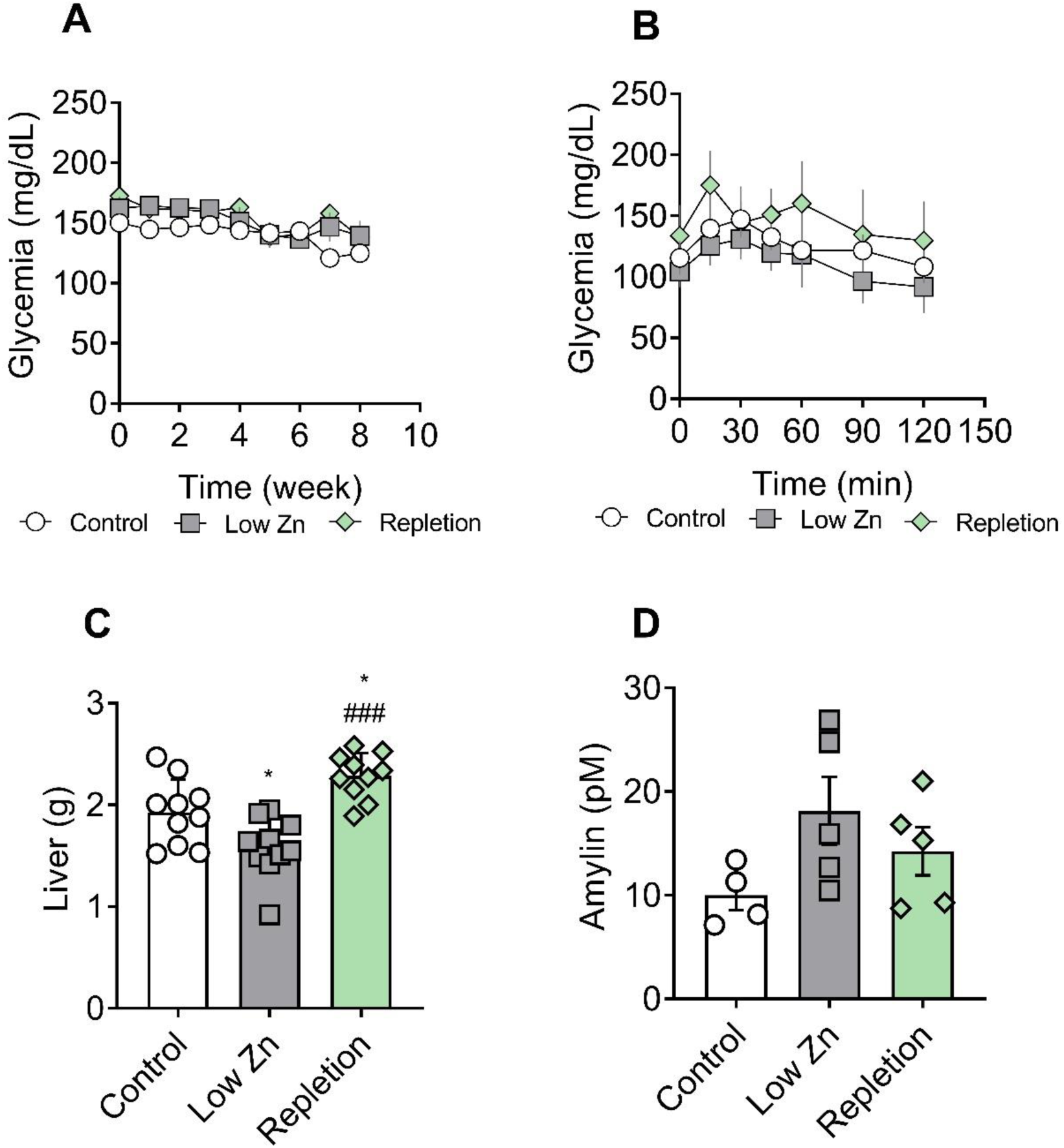
Effect of dietary zinc on metabolism. Capillary blood glucose without fasting weekly measurement (**A**) and oral glucose tolerance test performed in the 8^th^ week (**B**). Liver weight (**C**) and plasma amylin levels (**D**). Mean ± SEM, n=4 to 10. Two-way ANOVA followed by Tukey’s multiple comparison post-test (**A** and **B**) and one-way ANOVA followed by Tukey’s post-test (**C**). * p < 0.05 and ^###^ p < 0.001.

### Zinc restriction does not change the insulin-islet size relationship

Pancreas were fixed and embedded in paraffin, and sections were evaluated for insulin labelling by immunohistochemistry. All three groups showed a large distribution of islet sizes, spanning about 2 orders of magnitude in area (**Figure 5A, Figure D**: control; **Figure 5B, Figure E**: depletion; **Figure 5C, Figure F**: repletion). Insulin labeling was apparently more intense in smaller islets (**Figure 5G**). The integrated intensity over each islet showed a close similarity (**Figure 5H**), and thus the relation between insulin labeling and islet area presented a large distribution over orders of magnitude (**Figure 5I**).

**Figure 5.**
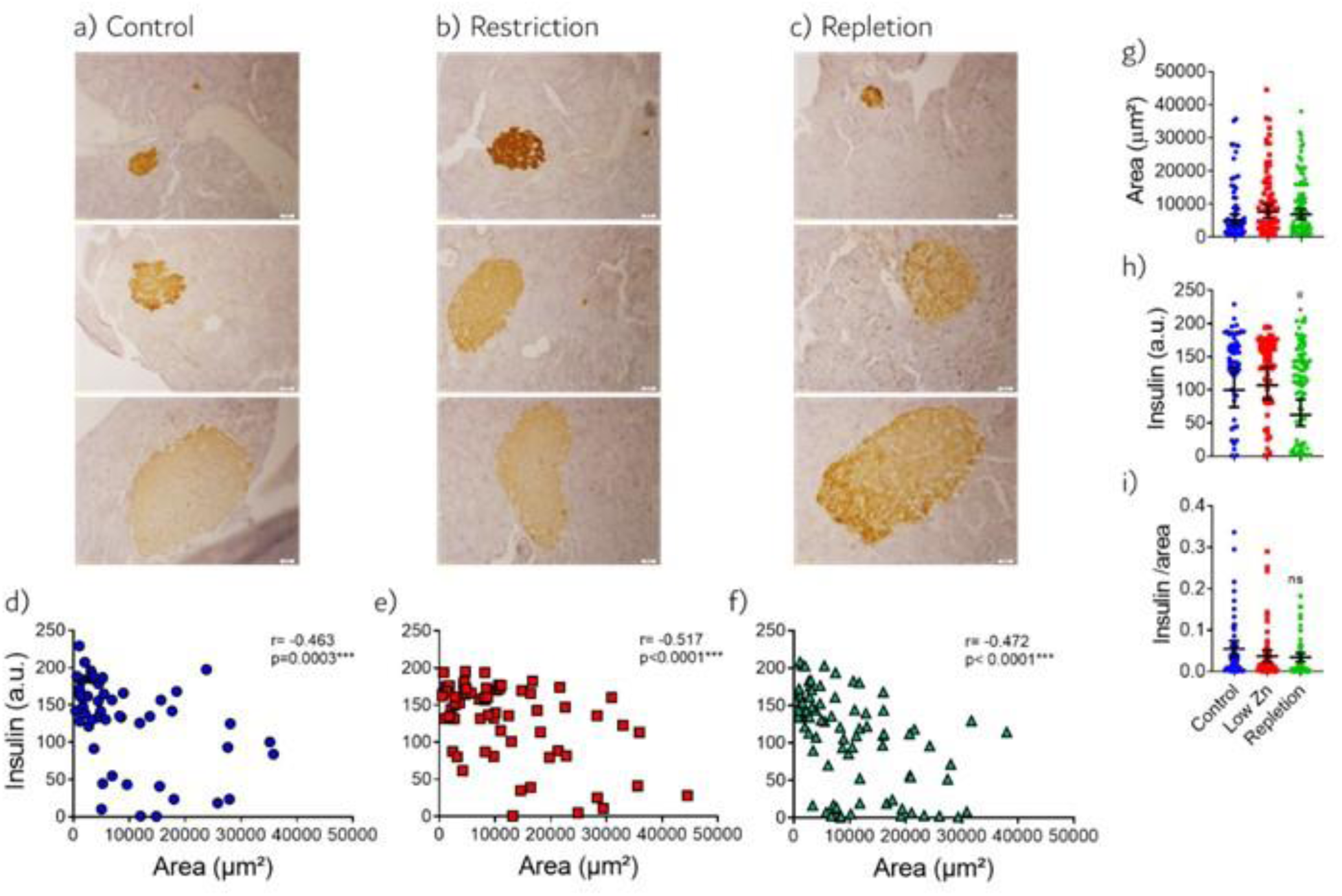
Representative images of the different dimensions of pancreatic islets. The islet area was evaluated by immunostaining for insulin in the control (**A**) Low Zn (**B**) and Repletion (**C**) groups. Semi-quantification analysis of islet insulin immunostaining per group, control (**D**), Low Zn (**E**) and Repletion (**F**). Comparison between groups of area quantification (µm^2^) (**G**). Comparison between groups of semiquantitative insulin immunostaining intensity (**H**). Relationship between insulin labeling intensity and islet area (**I**). n= 56-80 islets per group. Two-way ANOVA followed by Sidak’s multiple comparison post-test. * p < 0.05 vs control group; ^#^ p < 0.05 vs Low Zn group. Scale bar = 20µm, magnification 40x.

We observed a significative inverse relationship of insulin intensity and islet size (**Figure 5I**), indicating increased content in smaller islets. This relationship was higher in the depletion group (*Control*, r = -0.663, p = 0.0003; *Depletion*, r = -0.517, p < 0.0001) and reverted in the *repletion* group (r = -0.472, p < 0.0001).

Using bright field microscopy, no Congo red staining was observed in the islets of the three diet groups (**Figure 6**, upper slides), indicating absence of detectable amounts of amyloid deposits. This was confirmed by a lack of light-green birefringence in the islets when examining the same pancreatic tissue sections under polarized light (**Figure 6**, lower slides).

**Figure 6.**
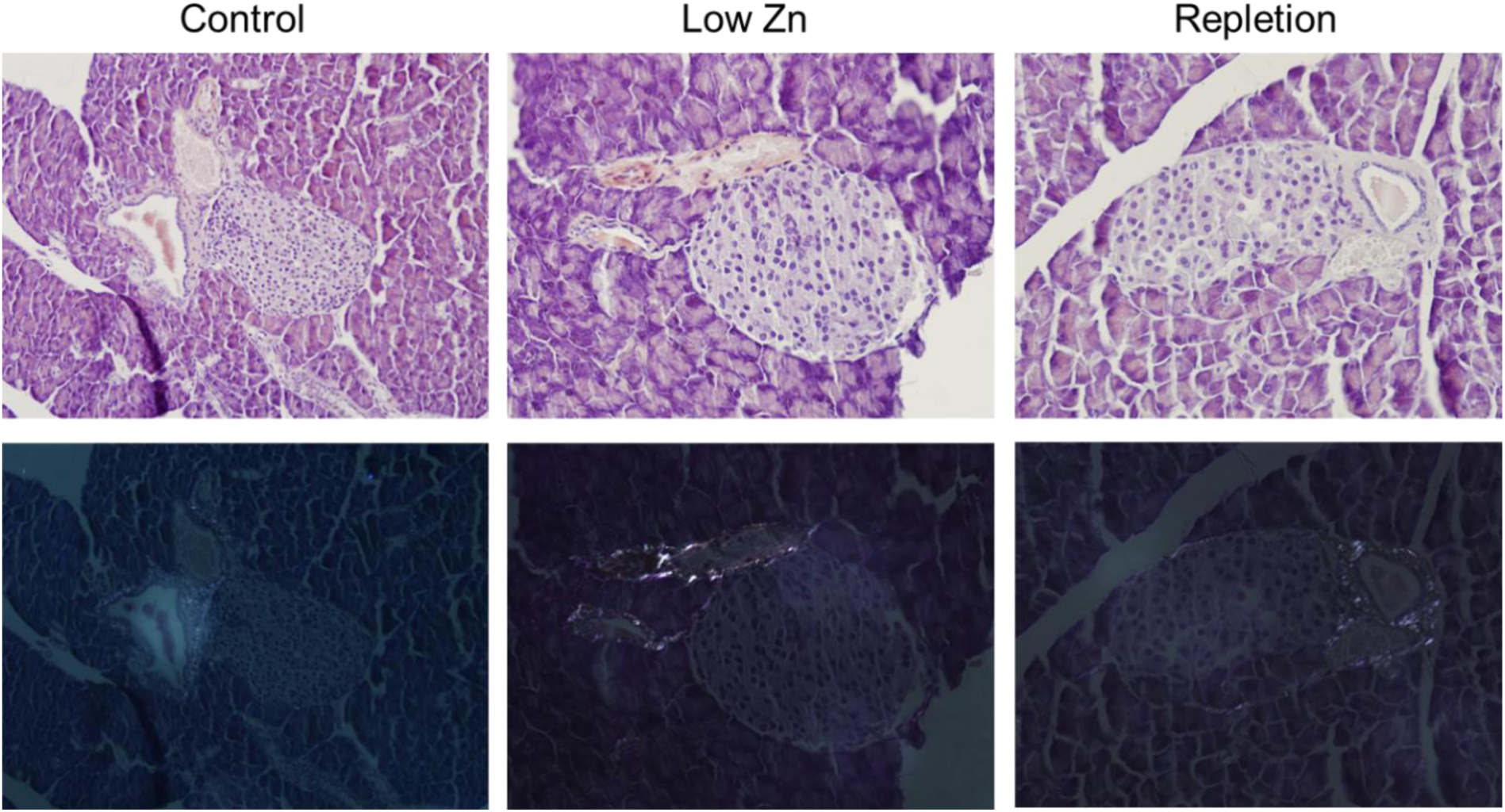
Lack of amyloid material in pancreas due to zinc dietary supply. Amyloid material was screened by Congo red staining of histological sections and representative islet images of the groups (n=3 animals/group) are visualized under bright field (upper slides) and polarized light (lower slides). Positive amyloid control (not show) was used to ensure validity of the method, and collagen birefringence around vascular system also indicates correct staining and lighting.

### Zinc restriction increases mechanical and thermal allodynia

The effect of zinc restriction and repletion on nociceptive response in adult mice was evaluated for both mechanical (von Frey) and thermal (Hargreaves) stimulus. The nociceptive response to mechanical stimuli was stable over the eight weeks experimentation for the *control* group, as expected for adult mice (**Figure 7A**). The zinc *restricted* and *repletion* groups showed progressive decrease in the stimuli threshold, compatible with increased nociceptive pain, a low withdrawal threshold plateau from the 4th week (p = 0.0045 restricted and p = 0.0044, *low zinc* vs *control group*) (**Figure 7B**). Zinc repletion by changing diets did not revert the mechanical allodynia within the following four weeks (**Figure 7B**).

**Figure 7.**
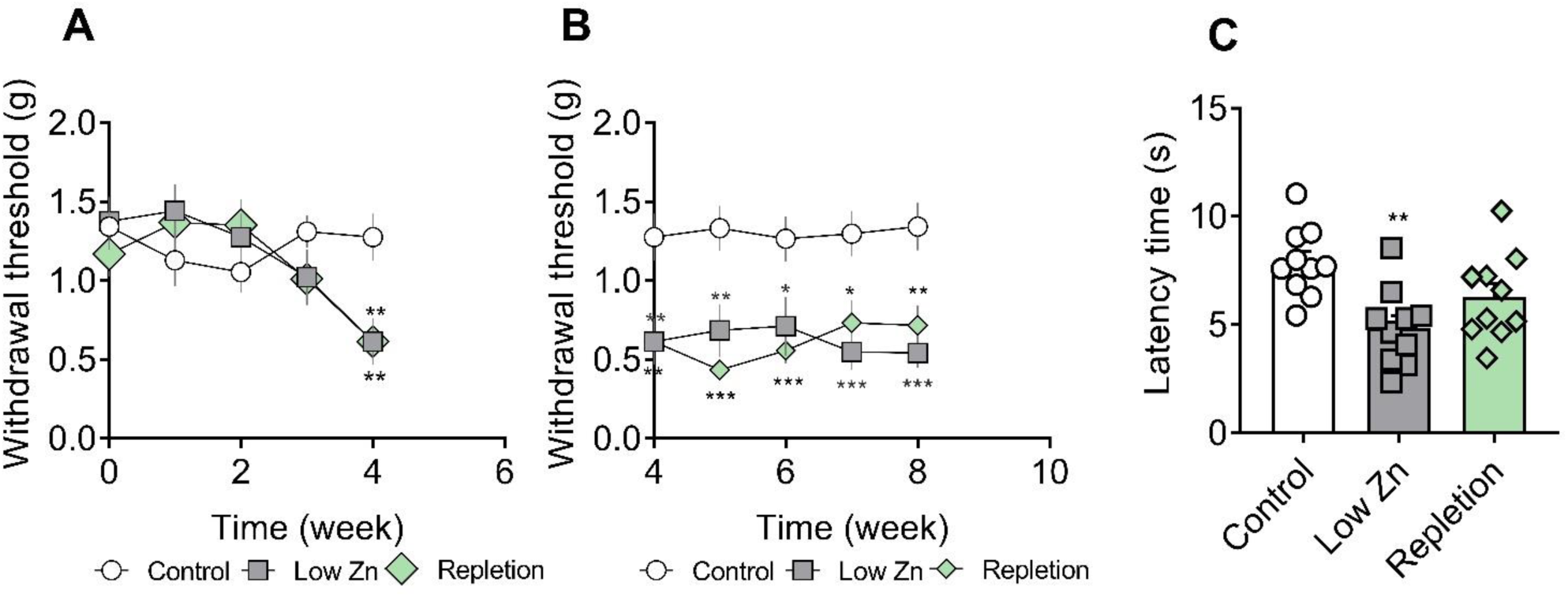
Effect of dietary zinc on mice mechanical and thermal allodynia. The mechanical nociceptive response was evaluated weekly in all experimental groups. Withdrawal threshold at first four weeks of zinc dietary restriction (**A**) and at four weeks after zinc reposition for the repletion group (**B**). The thermal sensitivity to heat was measured in the 8^th^ week of the protocol (**C**). Results are expressed as mean ± SEM (n=10-18). * p < 0.05 and ** p < 0.01 *** p < 0.001 compared to control group, Two-way ANOVA followed by Sidak’s multiple comparison post-test (**A** and **B**) and One-way ANOVA followed by Tukey’s multiple comparisons test (**C**).

The heat thermal allodynia was also evaluated. After eight weeks of intervention, the zinc *restricted* group showed a decreased latency time, compatible with greater sensitivity (p = 0.0025) **(Figure 7C).** The *zinc repletion* group was unable to change this response, with significant difference observed compared to *low zinc* group. Together, these data suggest that zinc restriction is independently sufficient to induce increased nociceptive pain in adult mice.

## 4. Discussion

Zinc deficiency is considered one of the main causes of morbidity in developing countries, one of the most prevalent nutritional deficiencies worldwide, and responsible for growth retardation in children. It is estimated that 17.3% of the population has an inadequate intake of zinc (Bailey et al. 2015).

In the present study with adult mice, we observed that the *control* group, supplied with normal content of zinc, showed an average body weight gain of 12%, while zinc *restriction* led to a 10% reduction in average total body weight at the end of the experimental protocol. Similarly, there are reports that post-weaning rodents that were subjected to a restricted zinc diet showed around 45% less body weight gain in Wistar rats and reduced weight gain and body growth in mice (Jing et al. 2015; Sisnande et al. 2020; Lima et al. 2023). These data demonstrate that zinc deficiency affects the body weight of mice at different stages of their development.

Despite affecting body weight gain, this could not be attributed to an unsatisfactory intake, since the experimental groups showed similar dietary intake. On the other hand, zinc repletion after its restriction appears to affect abdominal adipose tissue accumulation, showing an increase of 23% in total body weight, being significantly greater than the animals that remained under restriction and presented significant increase in abdominal adipose tissue. This effect was not observed in pigs when zinc supplementation was carried out aiming to improve weight gain, however without evidence of a prior restriction (Martinsson and Ekman 1976).

Thus, body weight, abdominal adipose tissue and liver data indicate that the zinc restriction – repletion cycle impacts overall metabolism in the adult mouse, without major impact on nutrient intake.

In adult mice, zinc restriction induced non-reversible nociceptive complications, despite being normoglycemic and without major alterations in insulin availability, suggesting that neurological misfunction may initiate before significative changes in glycemic manifestations. Nociceptive pain due to zinc restriction has been demonstrated previously by our group for weaning mice (Lima et al. 2023), which occurs along with degeneration of the endocrine pancreas (Sisnande et al. 2020). We believe that Dr. Bruce Ames’ Triage Theory applies in this case, which tells us that the organism responds differentially to micronutrient deficiency, and not equally in all parts, privileging essential functions for life (Ames 2006) although dictating evolution through nutrient-sensing mechanisms (Lima and Sisnande 2022). Consumption of the metal-chelating anti-nutrient phytate also results in allodynia (Matias et al. 2023). With this point of view, we understand that dietary zinc restriction compromised the allodynic response, while the endocrine activity of the pancreas would be largely maintained in the model and duration of the reported intervention. That way, these data indicate that zinc restriction is independently sufficient to induce increased nociceptive pain in adult mice.

We observed a negative dependence of insulin related to the islet size. This finding agrees with similar results from other research groups (Kim et al. 2009; Huang et al. 2011; Corbin et al. 2021). Our data demonstrated a significative decrease in insulin labeling in the repletion group, which seems to originate from larger islets with lower insulin labelling. However, hormone proteoforms with dissimilar processing stages can be found in islets (Sisnande et al. 2024), which cannot be inferred from the immunolabelling data. The magnitude of the effect seems small, which might be due to a zinc storage effect in adult animals. However, these results indicate direct effect of zinc restriction and repletion over the pancreatic β-cell health and islet function, related to insulin availability (production, secretion, degradation, and insulin proteoforms), necessary to maintain the glycemic control, which is in the core of diabetes definition.

Pancreatic islets in more than 90% of patients with type 2 diabetes present amyloid deposits (Mukherjee et al. 2015), which can be readily and specifically observed by Congo red staining, even without polarized light (Westermark et al. 2011). In our samples, neither bright-field red staining nor birefringence were visualized in the tissue sections, suggesting that a low zinc diet did not induce the formation of detectable amounts of murine amyloid *in vivo*, although murine amylin forms amyloid *in vitro* (Palmieri et al. 2013) and oligomers *in vivo* (Erthal et al. 2018).

It should be noted that the low Zn group had twice the concentration of plasma amylin compared to control animals, as shown by the present results. This is in line with previous findings by ourselves, in which the gene expressions of amylin and BACE2 were markedly enhanced in the pancreatic islets of mice fed a high-fat diet and with hyperglycemia (Cardoso et al. 2023). Therefore, absence of amyloid deposits in the islets occurred despite the greater bioavailability of fibril-forming amylin, both circulating and locally in the islet itself. It is important to notice that transgenic mice overexpressing human amylin form amyloid deposit *in vivo* in the pancreas as detected by ultrastructural techniques (Janson et al. 1996), and thus not necessarily the lack of Congo Red detection can be taken as absolute absence of amyloid in mice pancreas. These findings certainly warrant further research to explore in deeper detail a yet undiscovered role for zinc and other essential micronutrients in the neurodegeneration, chronic pain development and endocrine disruption at the perspective of the system biology and physiology.

In conclusion, nutritional zinc deficiency induces changes in mechanical and thermal nociceptive allodynia in adult mice, which indicates a drift in the neural physiology, which is not likely to be reversed under the conditions of the present study. Thus, a direct link between zinc deficiency and changes in nociceptive responses in adult animals are demonstrated. Further studies should investigate the mechanisms underlying this inflammatory panorama, the nociceptive pathways and global nutritional status.

## Acknowledgments

We would like to thank the staff of the Laboratory of Molecular Pharmacology – UFRJ (Prof. Jorge Luiz Mendonça Tributino and Prof. Newton Gonçalves de Castro) for access to their microscopy platform.

## Data Availability

The data generated during and/or analyzed during the current study are available from the corresponding author on reasonable request.

## Author Contributions

**DOM -** Conceptualization; Data curation; Formal analysis; Investigation; Methodology; Visualization; Writing - review & editing.

**BLRS** – Investigation; Methodology; Writing - review & editing.

**AFM** – Investigation; Methodology; Writing - review & editing.

**LPS** - Investigation; Methodology; Writing - review & editing.

**LMA** - Investigation; Methodology; Writing - review & editing

**LEMC** - Investigation; Methodology; Writing - review & editing

**ALPM -** Conceptualization; Data curation; Formal analysis; Methodology; Supervision; Visualization; Writing - review & editing.

**LMTRL -** Conceptualization; Data curation; Formal analysis; Funding acquisition; Investigation; Methodology; Project administration; Supervision; Visualization; Roles/Writing - original draft; Writing - review & editing.

